# Effects of longitudinal bending stiffness and midsole foam on running energetics

**DOI:** 10.1101/2024.05.07.592978

**Authors:** Damion Perry, Herlandt Lino, Montgomery Bertschy, Wouter Hoogkamer

## Abstract

**Background:** Advanced footwear technology has become commonplace in the competitive running world. However, a systematic exploration of the effects of each of the primary components has not yet been presented.

**Purpose:** To quantify running economy and step parameters in four different shoe conditions: PEBA shoes with plate, PEBA shoes without plate, EVA shoes with plate and EVA shoes without plate.

**Methods:** Participants ran at 14 km/h (n = 8 males) or 12 km/h (n = 6 females) on a treadmill. The shoe order was randomly assigned for the four shoe conditions, where the participants wore each shoe twice in a mirrored order.

**Results:** There was a significant reduction in metabolic power (1.0%) associated with the presence of a plate, as well as a similar reduction based on foam type (1.0%). However, there was no significant interaction between presence of plate and foam type. Adding a plate significantly reduced metabolic power for EVA by 1.3% but not for PEBA. PEBA foam significantly reduced metabolic power by 1.3% in shoes without a plate, but not for the shoes with a plate. Step frequency and contact time were similar between shoes and not correlated to running economy improvements.

**Conclusions:** Starting from a baseline condition with traditional foam without a plate, either adding a plate or using PEBA foam improved running economy with a similar amount (1.3%). Adding the alternate second feature non-significantly improved running economy with an additional 0.6%. The benefit of both technologies combined (1.9%) was smaller than the sum of its parts (1.3% each).

**Key points:**

- The use of a PEBA foam midsole or the addition of a carbon fiber plate improves running economy a similar amount when compared to a traditional EVA running shoe without a plate.
- Contrary to our hypothesis, the benefit of both technologies combined (1.9%) was smaller than the sum of its parts (1.3% each).

## 1 Introduction

The emergence of advanced footwear technology (AFT) has dramatically altered the world of distance running. AFT shoes combining lightweight, resilient midsole foams with rigid moderators and pronounced rocker profiles (Frederick, 2022), have been shown to drastically improve running economy and performance. For instance, the Nike Vaporfly 4% was so named after findings by Hoogkamer et al. (2018) that showed a 4% improvement in running economy (the rate of metabolic energy consumption (W/kg) at a defined submaximal running velocity), later repeated by Barnes and Kilding (2019) and Hunter et al. (2019). This 4% improvement in running economy has resulted in improvements in elite running performance in both men and women across all road running events (Bermon et al., 2021; Rodrigo Carranza, González-Mohíno, et al., 2023; Senefeld et al., 2021).

Intriguingly, while most running brands currently have AFT models on the market, there is still an ongoing debate about which AFT element is more important and how each improves running economy (Burns & Tam, 2020; Hébert-Losier & Pamment, 2023; Kram, 2022; Matijevich et al., 2023; Nigg et al., 2021; Patoz et al., 2022). The public debate seems to particularly focus on the use of the carbon fiber plates (Dyer, 2020; Muniz-Pardos et al., 2021), even though scientific studies suggest that the effects of carbon fiber plates on running economy are small (for review see: Ortega et al., 2021). Roy and Stefanyshyn (2006) were the first to report an improvement in running economy (0.8%) due to increased longitudinal bending stiffness from carbon fiber plates. Results from studies since then have been mixed. Studies using flat carbon fiber insoles have reported no significant differences in running economy (Beck et al., 2020; Cigoja et al. 2021; Flores et al., 2019) or even deterioration in running economy (Day & Hahn, 2020). Another study, using an embedded curved carbon fiber insole, observed running economy improvements, ranging from 1 to 4% at different speeds for groups with varying training backgrounds, in shoes with traditional midsole foam (ethylene vinyl acetate, EVA) (Rodrigo Carranza et al., 2023).

Numbers on the running economy effects of curved carbon fiber plates embedded in modern compliant and resilient midsole foams are less readily available. There are some indirect indications that the plates contribute less to the running economy improvements than the use of modern foam (Healey & Hoogkamer, 2022; Hoogkamer et al., 2019). By comparing running biomechanics in Nike Vaporfly 4% prototype shoes versus baseline racing flats, Hoogkamer et al. (2019) found that the plate reduced negative work in the metatarsal phalangeal joint compared to baseline racing flats (Hoogkamer et al., 2019). Quantifying the mechanical energy return from plate bending and midsole foam compression revealed that the mechanical energy return from plate bending was 50 times smaller than the mechanical energy return from foam compression (Hoogkamer et al., 2019). Furthermore, Healey and Hoogkamer (2022) made medio-lateral cuts through the midsole and plate in Nike Vaporfly 4% (AFT) shoes, effectively reducing the longitudinal bending stiffness. Compared to intact shoes, differences in running economy were small (~0.5%) and insignificant. Cutting the plate medio-laterally did not remove the plate and as such the plate could still fulfill other functions such as stabilizing the more compliant foam and spreading out the forces under foot over a larger area (Healey & Hoogkamer, 2022; Kram, 2022).

While there is accumulating evidence that increased midsole thickness does not improve running economy (Barrons et al., 2023; Bertschy et al., 2023), results on the independent effects of midsole foam compliance and resiliency are limited. Frederick et al. (1983) observed small (<1%) differences in running economy between shoes with different hardness EVA foams and butyl foam. Worobets et al. (2014) observed a 1% running economy improvement from EVA to more compliant and resilient TPU (thermoplastic polyurethane) midsole foam.

To fully understand the independent contributions of the plate and the foam, identical AFT shoes with and without plates, and with and without modern midsole foam should be compared. Therefore, the purpose of this study was to quantify how midsole foam type and presence of a curved carbon fiber plate independently affect running economy, by using shoes identical in geometry and mass, but differing in foam, with and without an embedded, curved carbon fiber plate. We quantified how highly compliant and resilient foam (polyether block amide, PEBA) affects metabolic rate compared to more traditional midsole foam (EVA), both with and without a curved carbon fiber plate embedded in the midsole. We anticipated that both switching from EVA to PEBA foam, and adding a carbon fiber plate would improve running economy, with the combined effect being larger than the sum of each separate effect (Kram, 2022). We hypothesized that running economy would be worst in EVA shoes without a plate and best in PEBA shoes with a plate, and that running economy in PEBA shoes without plate would be better than in EVA shoes with plate. We also hypothesized that step frequency would be slower (i.e., longer step length) and ground contact time longer in PEBA shoes with plate, than in the EVA shoes without plate.

## 2. Materials and Methods

### 2.1 Participants

We recruited 14 runners capable of running 5 km in 19 minutes (males) or 21 minutes (females) or an equivalent performance (e.g., 10 km in 39 minutes, marathon in 3:00 hours for males; 10km in 44 minutes, marathon in 3:22 hours for females), fitting men’s US9 or women’s US8 sized shoes. Exclusion criteria were a lower extremity injury in the past two months, surgery in the past twelve months or any existing orthopedic, cardiovascular, or neuromuscular conditions. All participants gave written informed consent that followed the guidelines of the University of Massachusetts, Amherst IRB (protocol # 2927).

### 2.2 Shoe conditions

We used the Saucony Endorphin Pro (Saucony, Waltham, MA, USA) which has a PEBA based midsole (“PWRRUN PB” bead foam) and an embedded curved carbon fiber plate as our AFT condition (PEBA with plate). Saucony provided three other custom versions: PEBA without plate - with the same foam, but without plate; EVA with plate - with EVA foam and an embedded curved carbon fiber plate; and EVA without plate - with EVA foam, without plate. All shoes were equalized for mass, matching the mass of the EVA with plate condition (Table 1), using adhesive steel weights along the base of shoe upper. Weights placed along the top of the shoe midsole along the base of the upper more closely replicate the weight distribution of the heavier shoe conditions, where the additional mass is due to the heavier midsole foam and midsole embedded carbon fiber plate.

**Table 1.** Shoe properties for the experimental shoe conditions (men’s US 9).

|  | EVA<br>without<br>plate | EVA<br>with plate | PEBA<br>without<br>plate | PEBA<br>with plate |
| --- | --- | --- | --- | --- |
| Mass (g) [before weight matching] | 228 | 248 | 196 | 215 |
| Compression Stiffness (N/mm) | 81.4 | 89.8 | 76.3 | 83.9 |
| Energy Return (%) | 75.2 | 80.4 | 83.6 | 87.7 |
| Longitudinal Bending Stiffness (Nm/rad)<br>(3-point) | 22.2 | 35.4 | 17.1 | 28.2 |
| Longitudinal Bending Stiffness (Nm/rad)<br>(Shoe Flexer) | 12.4 | 18.1 | 8.1 | 19.5 |

### 2.3 Mechanical testing

The shoes’ compression stiffness, energy return and longitudinal bending stiffness were measured using a material testing machine (Instron ElectroPuls 10000; Instron, Norwood, MA, USA) and a flex tester (Shoe Flexer; Exeter Research Inc., Brentwood, NH, USA). Midsole compression stiffness was measured by applying a parabolic force profile with a peak magnitude of 1800 N for 185 ms onto the heel of the shoe with a rounded probe (diameter = 43 mm, radius of curvature = 21.5 mm). Compression stiffness was calculated from the maximum force applied and the maximum displacement. Energy return was calculated from the area under the force-deformation curve during loading and unloading. Longitudinal bending stiffness was measured by two methods. A recently often used method is the 3-point bending test. For this the forefoot of the shoe was placed on two supports 80 mm apart. Midway between these supports, at the approximate location of the metatarsal phalangeal joint, a 7.5 mm displacement from a 50 N preload was applied for 20 cycles at 15 mm/s. Bending stiffness was calculated for the final three cycles from the applied force, displacement of the Instron tip, and the distance between the supports (Ortega et al., 2021). Since this test is susceptible to error due to foam deformation (Healey et al., 2021), we additionally measured the longitudinal bending stiffness using a standard flex tester, calculating stiffness for the final five of 50 30-degree flexion cycles.

### 2.4 Experimental protocol

Participants wore their own shoes for a warm-up period of at least 5 minutes at the test pace of 14 km/h (6:54-minute mile pace) (n = 8 males) or 12 km/h (8:02-minute mile pace) (n = 6 females), while breathing through a mouthpiece attached to an expired gas analysis system. After a five-minute rest period following the warm-up, participants completed eight, 5-minute trials at 14 km/h for males and 12 km/h for females on a force-measuring treadmill with a rigid deck (Treadmetrix, Park City, UT, USA). The shoe order was randomly assigned for the four shoe conditions, where the participants wore each shoe twice in a mirrored order (e.g., ABCDDCBA). Researchers removed and applied shoes between trials during the five-minute rest period to keep participants unaware of the shoe condition from handling the shoes, to reduce any associated personal bias. During the experimental running trials, the submaximal rates of oxygen uptake and carbon dioxide production were measured using a breath-by-breath expired gas analysis system (TrueOne 2400, ParvoMedics, Salt Lake City, UT, USA). The expired gas analysis system was calibrated right before testing and recalibrated midway through testing (i.e., after each condition was run in once) to reduce the effects of long-term analyzer drift. We quantified running economy as metabolic power based on the rates of oxygen uptake and carbon dioxide production over the last 2-minutes of each trial, using the Perronet and Massicotte (1991) equation and expressed metabolic power in W/kg. All participants maintained a respiratory exchange ratio <1.00 (0.81-0.94). Ground reaction force data was recorded at 1000 Hz during the last 30 s of each trial. Ground reaction force data was filtered using a dual-pass Butterworth filter with a 20 Hz cut-off (Healey & Hoogkamer, 2022; Hoogkamer et al., 2019). Touch down and toe off were determined using a 25 N vertical ground reaction force threshold to calculate ground contact time and step frequency.

### 2.5 Statistics

We present all results as mean ± SD values in the text and figures. We used linear mixed-effects models (LMEM) to evaluate the effects of foam type, plate presence, and their interaction on running economy, step frequency and contact time. We used a traditional level of significance (α = 0.05), with Tukey’s HSD test corrections in post hoc analyses. Two LMEMs were created for analyzing metabolic power. Since both trial number (p < 0.001) and sex (i.e., speed, p = 0.006) had significant effects, they were included in the models alongside with shoe (and foam/plate interaction) as fixed-effects, participant as a random intercept, and metabolic power as the output variable to test the effect of shoe and the foam/plate interaction, respectively. Two LMEMs were created for analyzing spatiotemporal variables step frequency and contact time. Shoe (and foam/plate interaction) was included as a fixed-effect, participant as a random intercept, and step frequency and contact time as the output variables, respectively. We performed linear regression analysis to assess potential correlations for between-shoe changes in running economy versus body mass, step frequency, contact time and between-shoe changes in step frequency and in contact time. We performed analyses in Python (Python Software foundation, https://www.python.org/) and R (R software Version 2022.12.0.353; The R Core Team, Vienna, Austria)).

## 3. Results

For running economy, there was a significant 0.97% reduction in metabolic power with the presence of the plate (13.64 ± 1.09 W/kg without plate to 13.51 ± 1.12 W/kg with plate; p < 0.001; ES = 0.12) as well as a significant 0.97% reduction based on foam type (13.64 ± 1.09 W/kg for EVA to 13.51 ± 1.11 W/kg for PEBA; p < 0.001; ES = 0.12; Figures 1, 2). However, there was no significant interaction between the presence of plate and foam type (p = 0.211). Running economy was worst in the EVA shoes without plate (13.73 ± 1.06 W/kg) and best in the PEBA shoes with plate (13.46 ± 1.12 W/kg, 1.93%; p < 0.001; ES = 0.24), but was similar in the PEBA shoes without plate and the EVA shoes with plate (0.00%, p = 1.0). Adding a plate significantly reduced metabolic power for EVA by 1.30% (p = 0.004, ES = 0.16), but not for PEBA (−0.64%, p = 0.325). PEBA foam resulted in 1.30% lower metabolic power for shoes without a plate (p = 0.004, ES = 0.16), but not for the shoes with a plate (−0.64%, p = 0.328).

**Figure 1.**
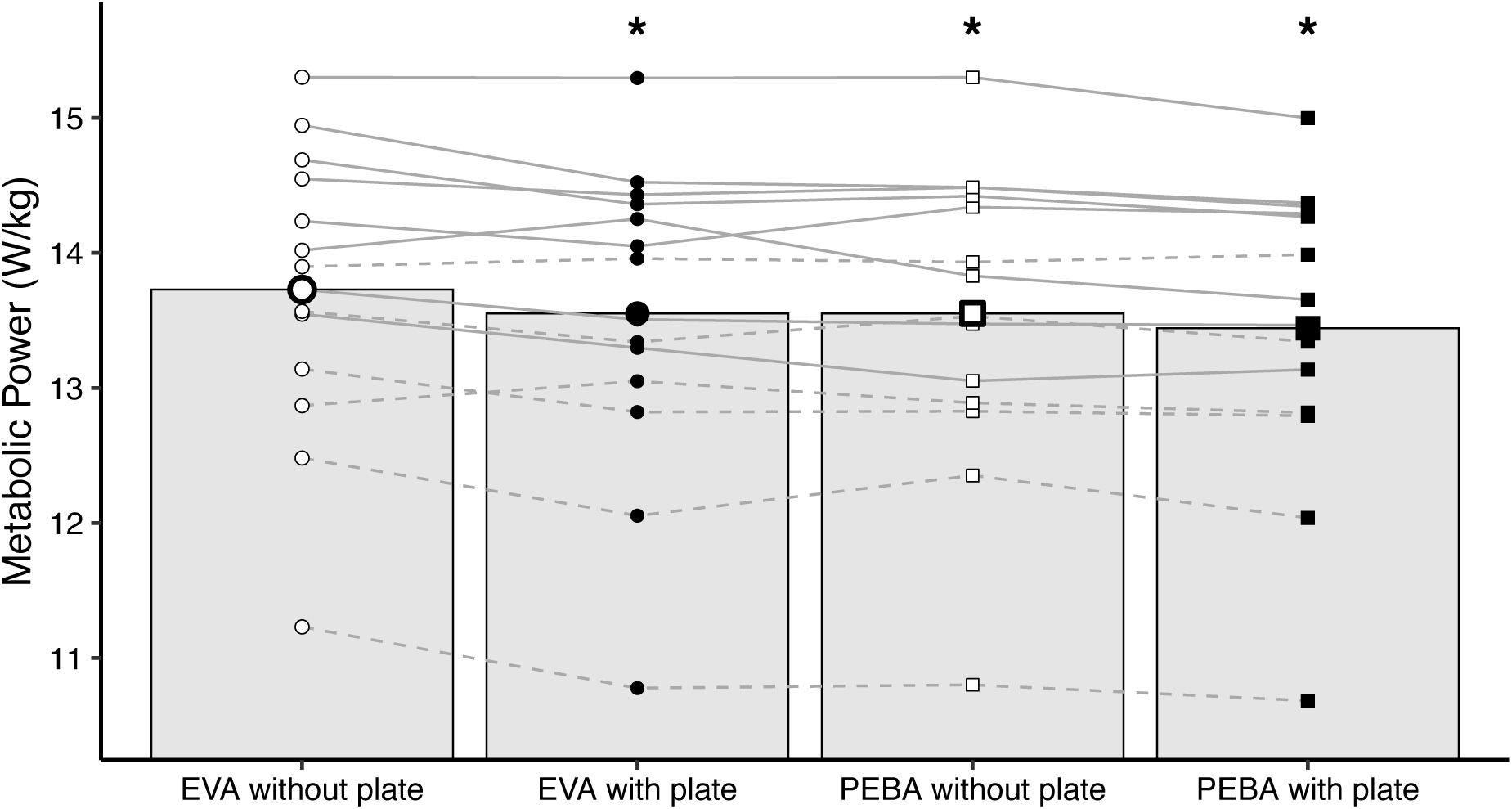
Running economy quantified as metabolic power in W/kg improved from EVA shoes without plate to PEBA shoes with plate. Black line represents average across all participants; grey lines represent individual participants’ average of two trials; solid lines represent male; dashed lines represent female; circles represent EVA; squares represent PEBA; open shapes represent without plate; closed shapes represent with plate.

**Figure 2.**
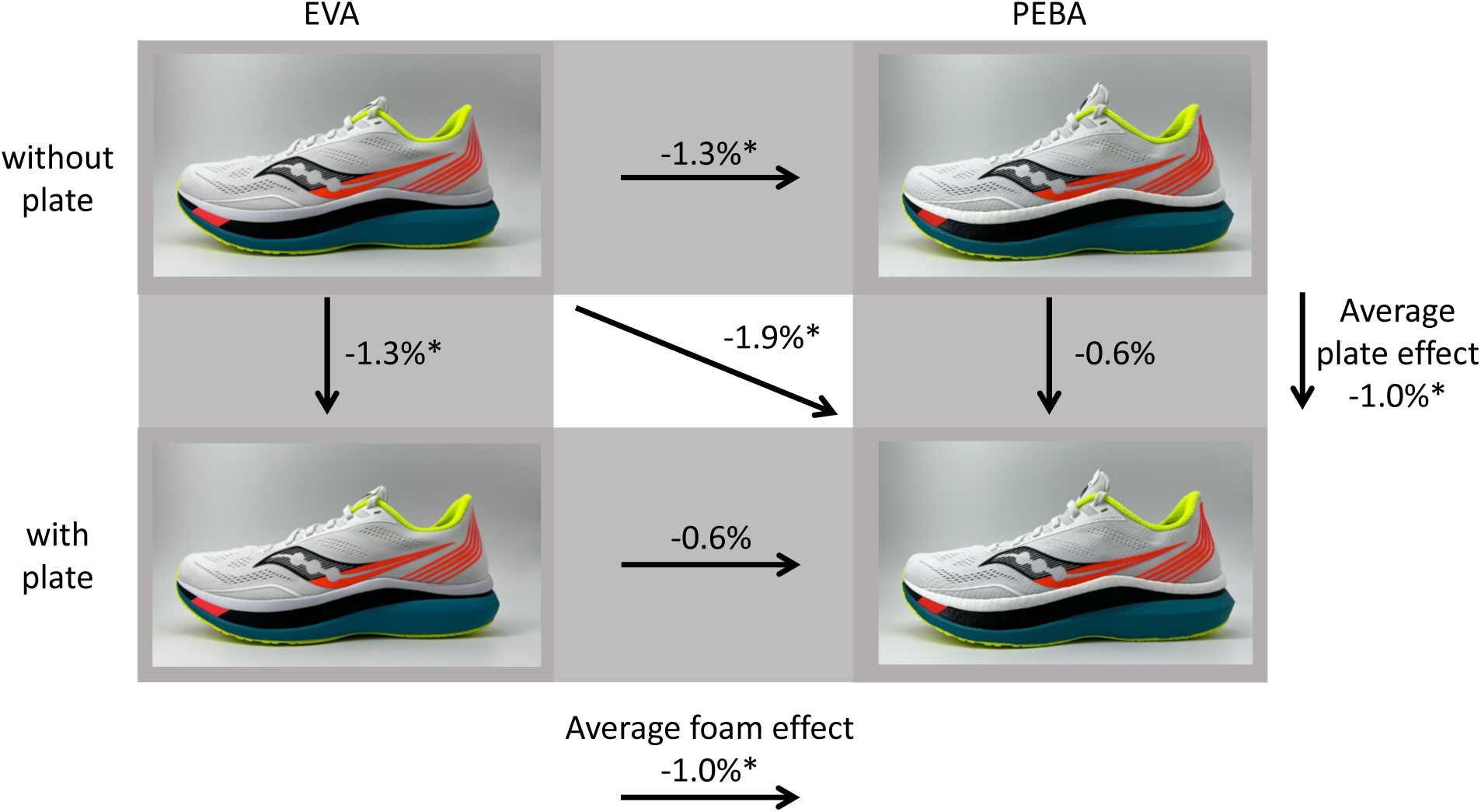
Running economy differences between shoe conditions for all participants. Shoes with and without a plate appear identical, foam differences between shoes are visible upon close inspection. * denotes statistical significance.

Step frequency and contact time were not different between shoes (Table 2). Further, improvements in running economy between shoes were not significantly correlated to the runners’ body mass, step frequency, contact time or to between-shoe changes in step frequency or in contact time (all r^2^ ≤ 0.17, p ≥ 0.145).

**Table 2.** Step frequency and contact time were similar between shoe conditions.

|  | EVA<br>without plate | EVA<br>with plate | PEBA<br>without plate | PEBA<br>with plate |
| --- | --- | --- | --- | --- |
| Metabolic Power (W/kg) | 13.73 ± 1.07 | 13.55 ± 1.14 | 13.55 ± 1.18 | 13.46 ± 1.12 |
| Step frequency (steps/min) | 175.6 ± 7.0 | 175.6 ± 6.7 | 175.6 ± 7.2 | 175.2 ± 6.9 |
| Contact time (s) | 0.216 ± 0.015 | 0.216 ± 0.014 | 0.216 ± 0.014 | 0.216 ± 0.015 |

## 4. Discussion

The purpose of this study was to quantify how midsole foam type and presence of a carbon fiber plate independently affect running economy, by using shoes identical in geometry and mass, but different in midsole foam type, with and without an embedded, curved carbon fiber plate. Confirming our first hypothesis, running economy was worst in EVA shoes without a plate and best in PEBA shoes with a plate. Running economy in PEBA shoes without a plate was similar to, not better than, in EVA shoes with plate. Starting from a baseline condition with traditional (EVA) foam without a plate, either adding a plate or using PEBA foam significantly improved running economy with a similar amount (1.3%; Figure 2). Adding the alternate second feature improved running economy with an additional 0.6%, non-significantly. Contrary to what we anticipated, the benefit of both technologies combined (1.9%) was not greater than the sum of its parts (1.3% each). Across foam conditions the average plate effect was 1.0% and across plate conditions the average foam effect was 1.0% (Figure 2).

To specifically quantify the mechanical effects of foam compliance and resilience, and of the plate, we choose to match shoe mass across the experimental conditions. EVA foam is generally heavier than PEBA foam (~30 g for a full midsole, Table 1), and carbon plates weigh ~20g (Table 1). Based on the 1% rule for shoe mass (Franz et al., 2012; Frederick et al., 1984) and our findings here, the improvements in running economy from the plate without weight-matching can be anticipated to be smaller, ~0.8% vs. 1.0% across foams. On the other hand, the improvements in running economy from PEBA foam without weight-matching can be anticipated to be ~1.3% vs. 1.0% across plate conditions. This suggests that for AFT shoes on the market, without artificial weight-matching, modern foam contributes more to improved running economy than the carbon fiber plate; by about ~0.5% for the shoes in this study.

Importantly, the running economy improvements we observed here from EVA shoes without plate to PEBA shoes with plate were less than the ~4% improvements observed for other AFT shoes as compared to traditional racing flats (Barnes & Kilding, 2019; Hoogkamer et al., 2018; Whiting et al., 2022). These larger improvements were relative to traditional racing flats which differed from the AFT shoes in multiple aspects, while in the current study all shoes featured an identical stack height and rocker geometry, taller and more pronounced than traditional racing flats. Increased stack height has been suggested to improve running economy (Burns & Tam, 2020), but experimental data suggests that increasing stack height by itself does not improve running economy (Barrons et al., 2023), and, depending on the midsole foam properties, might even worsen running economy (Bertschy et al., 2023). To the best of our knowledge the effect of rocker geometry on running economy has not been studied for AFT shoes, but Sobhani et al. (2014) observed a worse running economy in non-weight-matched traditional shoes with pronounced rocker geometry. The smaller differences in the current study might also be related to differences between AFT shoe models of different brands. Different models of different brands appear more or less effective in improving running economy (Joubert & Jones, 2022). Joubert and Jones (2022) observed that without any weight-matching the Nike Vaporfly Next% (i.e., the shoe model most similar to the AFT shoes improving running economy by 4% versus traditional Nike racing flats (Barnes & Kilding, 2019; Hoogkamer et al., 2018; Whiting et al., 2022)) improved running economy by 2.7% versus traditional Asics racing flats, while the Saucony Endorphin Pro (i.e., the shoes used in the current study) improved running economy by 1.5% in their study.

It is worth noting that the EVA used in this study appears more compliant and resilient than that in shoes tested in other studies (Table 1; Hoogkamer et al., 2018, Worobets et al., 2014). The larger compliance is likely related to the fact that we applied 1800 N over a smaller area (simulating heel strike) as compared to Hoogkamer et al. (2018) where 2000 N was applied over the full midsole area, and due to greater midsole thickness, as increased thickness for the same material will lead to a greater compliance. Whether the difference in loading area can also explain the higher resilience value in our study is less clear. If the EVA foam used in this study is indeed more resilient that may account for the smaller running economy differences presented in this study compared to previous findings.

Although we did not set out to evaluate sex differences in running economy changes between shoes, a preliminary, small sample (n = 8 males vs. n = 6 females) analysis revealed no significant differences in the response to the different shoes between males (running 14km/h) and females (running 12 km/h). Numerically, adding a plate improved running economy more in female runners than in male runners (0.9-1.5% vs. 0.5-1.1%, Table 3), and PEBA foam improved running economy more in male runners than in female runners (0.8-1.4% vs. 0.5-1.1%, Table 3). Whether these observations are robust or confounded by running speed differences should be addressed more carefully in future studies.

**Table 3.** Average running economy differences between shoe conditions for all participants.

| Sex | Metabolic Power Differences (%) |  |  |  |  |  |
| --- | --- | --- | --- | --- | --- | --- |
|  | without plate to with plate |  | EVA to PEBA |  | EVA without plate vs. PEBA | EVA with plate vs. PEBA |
|  | EVA | PEBA | without plate | with plate | with plate | without plate |
| Male | -1.32% | -0.33% | 0.73% | 1.72% | 0.39% | 2.07% |
| Male | -2.34% | -1.06% | -1.89% | -0.65% | -2.93% | 0.42% |
| Male | -1.86% | 0.65% | -3.63% | -1.21% | -3.02% | -1.85% |
| Male | -0.78% | -0.81% | -0.42% | -0.44% | -1.23% | 0.37% |
| Male | -0.04% | -1.97% | -0.01% | -1.94% | -1.98% | 0.03% |
| Male | -1.63% | -0.07% | -1.86% | -0.31% | -1.92% | -0.24% |
| Male | -2.81% | -0.99% | -3.06% | -1.24% | -4.02% | -0.25% |
| Male | 1.65% | 0.99% | -1.35% | -2.00% | -0.36% | -2.96% |
| Female | -1.65% | -1.34% | -0.27% | 0.03% | -1.60% | 1.42% |
| Female | -3.37% | -2.54% | -0.99% | -0.14% | -3.50% | 2.47% |
| Female | -2.44% | -0.27% | -2.38% | -0.18% | -2.64% | 0.06% |
| Female | -4.09% | -1.09% | -3.82% | -0.80% | -4.87% | 0.29% |
| Female | 1.41% | -0.57% | 0.16% | -1.78% | -0.41% | -1.22% |
| Female | 0.43% | 0.40% | 0.31% | 0.19% | 0.64% | -0.17% |
| Difference of the means | -1.30% | -0.64% | -1.29% | -0.64% | -1.93% | 0.00% |
| Male | -1.14% | -0.48% | -1.42% | -0.77% | -1.90% | -0.29% |
| Female | -1.53% | -0.88% | -1.10% | -0.45% | -1.97% | 0.44% |

Contrary to our second hypothesis step frequency and contact time were not different between shoes. When comparing AFT shoes to traditional racing flats, slightly slower step frequencies (~1% or less; i.e., longer step lengths) and slightly longer contact times (~1% or less) are often, but not always, observed (Barnes & Kilding, 2019; Hoogkamer et al., 2018; Hunter et al., 2019; Joubert & Jones, 2022). Differences in shoe properties in the current study were not pronounced enough to elicit significant differences. Further, improvements in running economy between shoes were not significantly correlated to the runners’ body mass, step frequency, contact time or to between-shoe changes in step frequency or in contact time. All the evaluated between-shoe associations for change in running economy and change in step frequency or contact time had a r^2^ ≤ 0.17 and p ≥ 0.145.

### Limitations

To quantify how midsole foam type affects running economy, we compared a traditional EVA foam versus a modern PEBA foam. Importantly, foam properties such as compliance, hardness and resilience can differ substantially even within a specific midsole foam type, depending on the exact chemical composition and production process. Similarly, to quantify how plate presence affects running economy we compared shoes with and without a carbon fiber plate, but plate geometry and bending stiffness can vary substantially. As such, our results are specific to the foams and plates we used, and should be generalized with care to shoe models with other foam and plate properties, even if the generic midsole foam type is the same. The findings by Joubert and Jones (2022) highlight that not all AFT shoes improve running economy similarly. Combining our observed improvements with the study of Flores et al. (2019) who did not find differences in running economy from increased energy return midsole foam or increased longitudinal bending stiffness, confirms that not all foams and plates are similarly effective and that factors as foam compliance or hardness and plate geometry and stiffness matter. Participants ran at 14 km/h (males) or 12 km/h (females) in our study, and even though Hoogkamer et al. (2018) and Barnes and Kilding (2019) observed consistent running economy improvements across speeds ranging from 14 to 18 km/h, it is possible that the relative importance of foam versus plate changes at different speeds (Joubert et al., 2023).

### Conclusions

In this study, we quantified the effects of the most discussed AFT features (midsole foam type and presence of a plate) on running economy by using identical shoe models and weight-matching. Starting from a baseline condition of traditional EVA foam, adding a plate or using PEBA foam improved running economy by a similar amount (1.3%). Adding the alternate second feature improved running economy by an additional, non-significant 0.6%. The benefit of both technologies combined (1.9%) was smaller than the sum of its parts (1.3% each). Without the weight-matching between shoe conditions applied in this study, the improvements in running economy can be anticipated to be 0.3% larger from PEBA foam and 0.2% smaller from the plate. Although the public debate regarding AFT shoes in the market has targeted the use of carbon fiber plates, our findings indicate that modern midsole foam technology contributes more to improved running economy than carbon fiber plates.

## Acknowledgements

We thank John Kuzmeski and Zach Barrons for performing the compression testing. We thank Katherine Boyer, Joseph Hamill, Alec Jessiman, Andrea Paulson, Cory Hofmann and James Allen for fruitful discussions

## Authors’ contributions

Conceptualization: Wouter Hoogkamer; Experimental design: Damion Perry, Wouter Hoogkamer; Data collection: Damion Perry, Herlandt Lino, Montgomery Bertschy; Data analysis: Damion Perry, Herlandt Lino, Montgomery Bertschy, Wouter Hoogkamer; Writing - original draft preparation: Damion Perry, Montgomery Bertschy, Wouter Hoogkamer; Writing - review and editing: Damion Perry, Herlandt Lino, Montgomery Bertschy, Wouter Hoogkamer

## Declarations

### Compliance with Ethical Standards

#### Ethical approval

The study was performed in accordance with the ethical standards of the Declaration of Helsinki. Ethics approval was obtained from the University of Massachusetts Institutional Review Board (Protocol # 2927)

#### Informed consent

Written informed consent was obtained from all individual participants included in the study.

#### Funding

This research was partly supported by a research contract from Wolverine Inc. (Saucony) with the University of Massachusetts, Amherst

### Data availability

All data discussed in this manuscript are provided in the tables.

